# Reconfigurations in brain networks upon awakening from slow wave sleep: Interventions and implications in neural communication

**DOI:** 10.1101/2021.12.07.471633

**Authors:** Cassie J. Hilditch, Kanika Bansal, Ravi Chachad, Lily R. Wong, Nicholas G. Bathurst, Nathan H. Feick, Amanda Santamaria, Nita L. Shattuck, Javier O. Garcia, Erin E. Flynn-Evans

## Abstract

Sleep inertia is the brief period of impaired alertness and performance experienced immediately after waking. While the neurobehavioral symptoms of sleep inertia are well-described, less is known about the neural mechanisms underlying this phenomenon. A better understanding of the neural processes during sleep inertia may offer insight into the cognitive impairments observed and the awakening process generally. We observed brain activity following abrupt awakening from slow wave sleep during the biological night. Using electroencephalography (EEG) and a network science approach, we evaluated power, clustering coefficient, and path length across frequency bands under both a control condition and a blue-enriched light intervention condition in a within-subject design. We found that under control conditions, the awakening brain is typified by an immediate reduction in global theta, alpha, and beta power. Simultaneously, we observed a decrease in the clustering coefficient and an increase in path length within the delta band. Exposure to blue-enriched light immediately after awakening ameliorated these changes, but only for clustering. Our results suggest that long-range network communication within the brain is crucial to the waking process and that the brain may prioritize these long-range connections during this transitional state. Our study highlights a novel neurophysiological signature of the awakening brain and provides a potential mechanistic explanation for the effect of light in improving performance after waking.

**One sentence summary:** Blue-enriched light partially accelerates the rapid prioritization of long-range communication within the human brain that characterizes sleep inertia

## Introduction

Immediately after waking from sleep there is a temporary period of reduced alertness and performance. The impact of this so-called *sleep inertia* on behavioral performance measurements has been well-described, including impaired reaction times,^1,2^ memory,^3,4^ decision-making,^5,6^ and a variety of other cognitive functions.^7^ These behaviors are also associated with a perceived state of sleepiness,^4,7^ disorientation,^8^ poor mood,^9^ and misperceptions of performance.^1^

Existing sleep inertia research has associated the waking process with several neural changes that include increased delta power over posterior regions of the brain,^10,11^ reduced beta power across all brain regions,^11^ and increased functional connectivity of the default mode network.^12^ Interestingly, the links between observed impaired performance and the neural behavior during sleep inertia are also commonly associated with sleep-related neural processes and states of sleepiness due to homeostatic and circadian pressures.^13^ These observations suggest a complex orchestration of neural elements supporting the transition from sleep to wakefulness spanning these oppositional constructs. Preliminary research investigating this complexity in neural network changes has suggested broad functional connectivity changes post-sleep, with the default mode network and the delta and beta bands playing a critical role in the network changes transitioning from sleep to wakefulness.^12,14^ These connectivity changes, however, have yet to be characterized and it is unknown whether an intervention may moderate these brain changes.

Understanding how heterogeneous neural elements of the brain coalesce to produce behavior and subjective experience may be understood via a graph theoretical framework. Using this framework, the brain is visualized as a graph or network made up of a collection of nodes (specified brain regions) and edges (connections) that represent brain elements and the corresponding statistical relationship between them. Topological description of brain networks within this framework can provide quantitative insights into the underlying mechanisms that give rise to emerging neural properties such as: specialization and efficiency of information processing;^15^ a variety of cognitive phenomena;^15-17^ and abnormalities in neurological disorders.^18^ Two common metrics, *clustering coefficient* and *path length*, have been used to describe properties of many complex systems, from biological phenomena^19^ to higher level systems.^20^ *Clustering coefficient* estimates the tendency of a node’s neighbors within a network to also be linked. *Path length*, on the other hand, estimates the number of edges, on average, that must be traversed to connect any two nodes within a network. When these metrics are at intermediate levels, associated with neither random nor regular networks, they describe the properties of a *small-world network*, which has been established as a popular model to describe functional brain networks by facilitating both localized and distributed processing of information.

In the current study, we describe the neurophysiological profile of the awakening brain under ecologically relevant sleep and circadian pressures using graph theoretical analysis of functional connectivity with high density electroencephalography (EEG). Using clustering and path length, we see that sleep inertia is characterized by a global shift in these metrics immediately after waking. Moreover, we confirm, using a within-subject, randomized, crossover intervention design, that blue-enriched light exposure alleviates this sleep inertia effect. We interpret our findings within the context of discontinuity of neural elements and efficiency of brain processes while the brain transitions from sleep to wakefulness.

## Results

Electroencephalography (EEG) from eleven participants (6 female; 23.1 ± 4.4 y, range 19-35 y, see Supplemental Table S1 for more demographic information) recorded during nocturnal awakenings from slow wave sleep (SWS) were analyzed. Briefly, due to the functional associations with defined categories of oscillations within the brain^21^ and previous associations of band-specific spectral power and sleep inertia,^10-12^ we first assessed the evolution of *global* spectral power (i.e., average across all channels) of the EEG recorded during the four separate test bouts and compared them to the pre-sleep baseline assessment. Then, via pairwise connectivity (wPLI) to estimate the phase-based relationship between channels, we estimated graph metrics that purposefully targeted the segregating and integrating aspects of the network. The temporal evolution of these estimates were then assessed to evaluate the duration of sleep inertia. Previous studies assessing the impact of sleep inertia typically report severe behavioral impairments that resolve within 15-30 minutes of waking.^22^ Remaining mild impairment may take up to an hour or more to dissipate, depending on conditions such as sleep pressure, sleep stage at waking, and time of day.^22,23^ Given these previous findings and our behavioral results under similar conditions,^1,9^ we consider a neural change to be due to *sleep inertia* when, compared to baseline, there is a significant change in the assessed metric at the first test bout (i.e., 2 minutes after waking; Figure 1, BL vs T1_C_).

**Figure 1.**
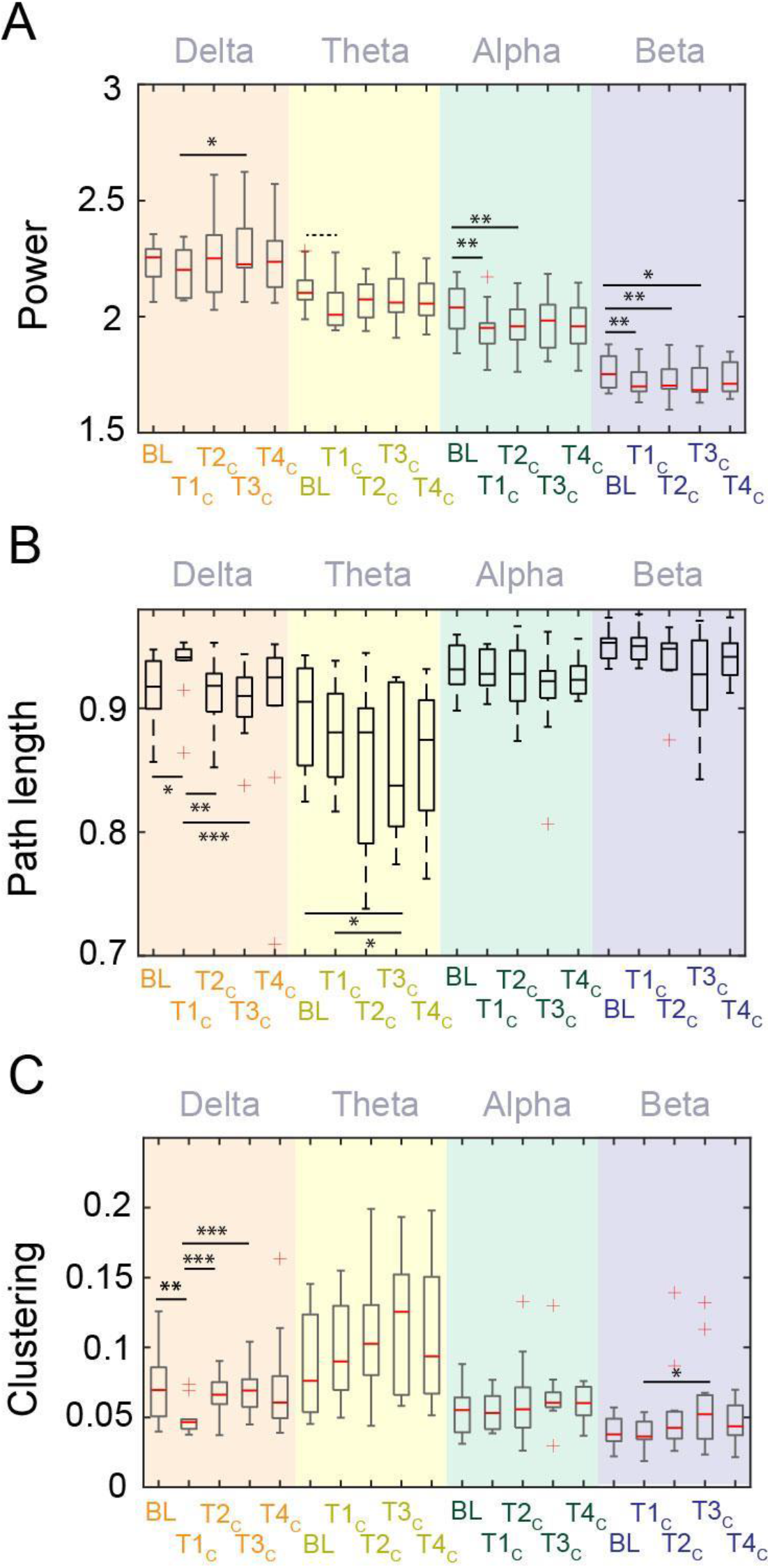
(A) Comparison for power, and (B) - (C) brain network properties across test bouts for each frequency band in the control condition (dim, red light). BL = baseline, T#_C_ = Test bout # during the control condition. \**p* < .05; \*\**p* < .01; \*\*\**p* < .001. Dashed line denotes marginally significant difference (*p* = .053).

### Global power of lower frequencies recovers faster than higher frequencies during sleep inertia

Figure 2A displays the average spectral power across all channels as a function of test bout. Statistical comparisons between these test bouts within each frequency band describe a complex coordination of neural firing that may be initially disturbed upon waking but then gradually recovers. Compared to participants’ global beta power prior to sleep (following moderate sleep restriction), beta global power was significantly reduced at T1_C_ (*t*(10) = 3.47, *p* = .006), T2_C_ (*t*(10) = 3.54, *p* = .005), and T3_C_ (*t*(10) = 2.98, *p* = .014). Similarly, compared to baseline, global alpha power was significantly reduced in the first two test bouts in the control (dim red light) condition (T1_C_: *t*(10) = 3.84, *p* = .003; T2_C_: *t*(10) = 3.56, *p* = .005). In the theta band, compared to baseline, we observed that the global power was marginally lower only at T1_C_ (*t*(10) = 2.2, *p* = .053). There were no significant differences between baseline and post-awakening test bouts within the delta band (all *p* > .05).

**Figure 2.**
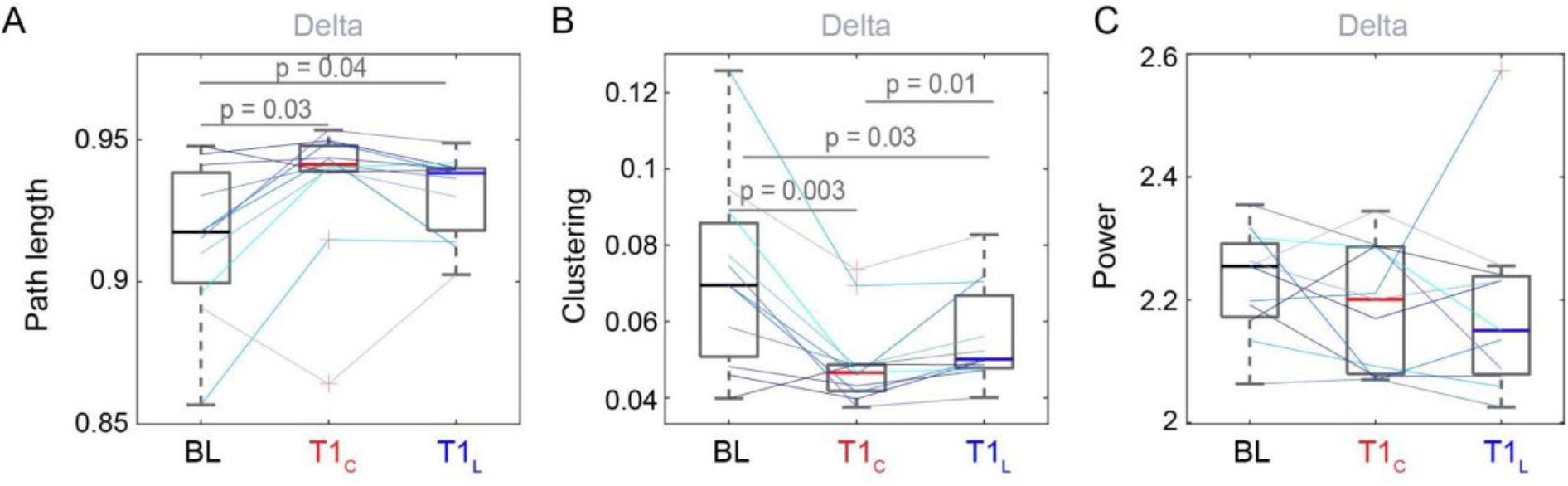
(A) - (B) Brain network properties comparing pre-sleep baseline (BL), control at T1 (T1_C_), and light at T1 (T1_L_) for the delta frequency band; (C) Similar comparison for delta power. Colored lines represent individual participants.

### Network properties of delta band connectivity display unique characteristics during sleep inertia

Next, we considered the global graph metrics of average shortest path length as a measure of integration and communication efficiency, in addition to average clustering coefficient as a metric of segregation. Under control conditions, path length was significantly increased in the delta band immediately after awakening compared to the pre-sleep baseline (*t*(10) = -2.52, *p* = .03). Path length reached baseline levels at T2_C_ (Figure 2B). Similarly, we observed a significant reduction of clustering coefficient immediately after awakening from SWS at night (T1_C_) compared to pre-sleep baseline (*t*(10) = 4.0, *p* = .002; Figure 2C). The clustering coefficient also returned to baseline levels at T2_C_ and persisted at that level. Except for a single effect of path length in the theta band significantly decreasing more than 30 minutes after awakening (*t*(10) = 2.65, *p* = .02), there were no significant differences in these metrics compared to baseline in other frequency bands. Thus, path length and clustering coefficient within the delta band reflect a robust sleep inertia signal, with an initial significant change immediately after waking, followed by recovery towards baseline at later time points.

### Blue-enriched light exposure at awakening attenuates neural network changes associated with sleep inertia in the delta band

We also assessed the effect of blue-enriched light as a way to potentially alleviate the neural changes associated with sleep inertia. Indeed, previous studies have shown that blue-enriched light has acute alerting properties, especially at night,^24,25^ and this effect has been demonstrated during the sleep inertia period.^9^ Figure 2 displays the effect of blue-enriched light on sleep inertia for each of the estimated global metrics across the frequency bands of interest for the baseline condition, the control condition at T1 (T1_C_), and the light intervention condition also at T1 (T1_L_). Importantly, we were interested in the effects that showed a difference between the control condition and the light intervention condition at this first time point at which the sleep inertia process is maximally influencing neural networks and behavior. See Supplemental Figure S1 for a visualization of all time points under the light condition.

Under the blue-enriched light intervention condition, path length at T1 in the delta band was not significantly different from control (*p* > .05). The clustering coefficient at T1 in the delta band, on the other hand, significantly increased in the light condition compared to control (*t*(10) = -3.04, *p* = .01); however, the clustering coefficient was still significantly lower than baseline at T1 in the light condition (*t*(10) = 2.56, *p* = .03; Figure 2B; see Supplemental Figure S2 for other frequency bands). Individual lines show a similar pattern for most participants. The clustering coefficient recovered to baseline levels at T2 in both conditions (see Supplemental Figure S3). Just as delta was the only frequency band reflecting sleep inertia under control conditions, the changes observed for delta path length and clustering under the light condition were not observed in other frequency bands, indicating a frequency-specific role of delta following awakening from SWS and in response to blue-enriched light.

There were no significant changes in power between the control and blue-enriched light conditions for any frequency band, suggesting that blue-enriched light does not attenuate changes in power immediately after awakening from SWS (Figure 2C).

### Sleep inertia is characterized by a global reduction in clustering and region-specific rescue with light

To understand regional contributions to the changes in the clustering coefficient, we next compared each electrode’s clustering coefficient between the baseline, control, and light conditions. Compared to the pre-sleep baseline, in the control condition all but one electrode showed a significant reduction in the clustering coefficient immediately after awakening (*q* < .05 after correction for multiple comparisons; Figure 3A). When exposed to light immediately after awakening, there was a trend for reduced clustering across the midline regions of the scalp (Figure 3B), but this did not survive correction for multiple comparisons (*q* > .05). When comparing the control and light conditions immediately after awakening (at T1), significant differences were observed in the right hemisphere, with higher clustering in the light intervention condition (Electrode F8: *t*(10) = -3.91; Electrode T8: *t*(10) = -3.95). We also inspected the regional differences in power and degree at each electrode (see Supplemental Figure S4).

**Figure 3.**
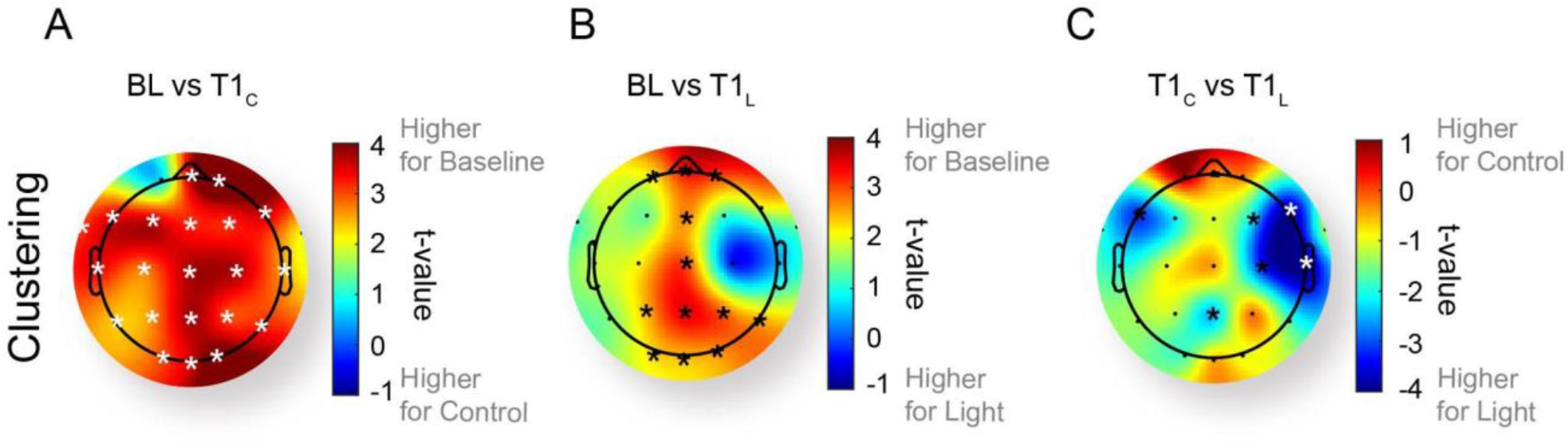
Change in clustering between (A) baseline (BL) and control at T1 (T1_C_), (B) baseline and light at T1 (T1_L_), and (C) control and light at T1 across scalp regions. White asterisks represent electrodes with significant difference on a paired-t test (*q* < .05) when corrected for multiple comparisons; black asterisks represent uncorrected significance (*p* < .05).

## Discussion

This is the first study to describe the changes to small-world network dynamics in the waking brain and the impact of blue-enriched light on this profile. Our analyses revealed significant reductions in clustering and increase in path length in the delta band immediately after awakening from SWS at night. Exposure to blue-enriched light attenuated the changes observed in clustering, suggesting a specific regime of segregation and integration driving the waking process. Network analyses, together with our investigation of an intervention, provide unique insight into the neurophysiological profile of sleep inertia beyond previous approaches.

### Small-worldness is disrupted during sleep inertia

Studies of characteristically small-world activity have shown that small-world properties are higher during sleep, especially in the slower oscillatory schemes, compared to wakefulness.^26,27^ Given this finding, we might expect higher small-worldness features immediately after waking from sleep if there is a slow transition from sleep to wakefulness. We observed, however, a significant reduction in clustering coefficient and increased path length immediately following sleep compared to waking levels. This change suggests a distinctive scheme towards a more random network with both an observed lower clustering of channels and simultaneously more segregation across them, indicating a decrease in small-worldness. Moreover, with fMRI-estimated connectivity, Liu and colleagues^28^ observed that under conditions of sleep deprivation, the estimated small-worldness increased relative to rested wakefulness, suggesting the brain may compensate for sleep deprivation via this mechanism. These findings may appear contradictory to what we observed in this study, but important experimental manipulations may provide a key understanding to how small-worldness shapes neural behavior during the critically important state of sleep, including how engaged participants are during a task, and the targeted network, as defined by nuances of the methodology or analysis (e.g., fMRI BOLD vs EEG power or EEG oscillatory scheme).

Interestingly, with EEG, Koenis and colleagues^29^ observed a decrease in small-worldness within the alpha band while awake but sleep-deprived, but reported no changes in the delta band. Importantly, these findings were reported while participants were engaged in a reaction time task. Given the similarities with our study design, the differences in our outcomes may reflect a unique profile of the waking brain that is distinct from the state of sleep deprivation itself. It is notable that immediately after waking, the primary features of small-worldness describe a unique aspect of brain segregation and integration, independent of both sleep-like features and the influence of sleep deprivation. This finding may indicate that, while participants are engaged in a task, sleep inertia is characterized by neural network reconfigurations which arise due to the disruption of SWS into an awake state. These reconfigurations place the brain network into a state of high segregation. Future research is needed to explore these findings.

### Global power suggests a prioritization scheme may underlie sleep inertia

Previous research seeking to describe the waking brain has focused on EEG power. Oscillations within the brain characterize rhythmic activity of subpopulations of neurons. It has been observed that specific cognitive functions are associated with different oscillatory schemes,^21,30,31^ where slower frequencies have been associated with top-down processes like executive actions and attention,^32,33^ higher frequencies are associated with motor control or maintenance of sensorimotor behaviors,^34^ and even higher frequencies with localized, specific, and rapid computations.^35^ Studies of sleep inertia have observed the waking brain exhibiting “sleep-like” attributes like high delta power and low beta power, compared to rested wakefulness.^10,11,36^ Notably, our baseline testing was conducted following mild sleep restriction, which may have elevated delta power in our baseline measures relative to rested wakefulness,^37^ and thus dampened our ability to detect a difference between pre-sleep and post-sleep delta power. This suggests, however, that delta power itself is not a unique signature of the waking brain. Further, we extended observations of EEG power to show the relative rate of recovery of these frequencies beyond timepoints previously tested (i.e., beyond 10-25 minutes post-sleep). We observed a marginally significant difference in the global power between baseline and the control condition in the theta band approximately 2 minutes after waking up and significant differences in global power between baseline and the control condition up to approximately 17 minutes in the alpha band, and approximately 32 minutes in the beta band after waking up. While none of these observations were different from each other across post-wake test bouts, this finding suggests that there is a measurable change in global power in these bands that is sustained for different lengths of time. For example, the recovery to baseline was faster for theta frequencies and longest for beta power, taking at least 30 minutes to return to pre-sleep levels. These observations support and extend findings from others who also reported reductions in alpha and beta activity immediately following awakening,^10,11,36^ but did not report the subsequent time course of recovery.

Our observations, overall, suggest that the broader organization of the brain may underlie the slower dissipation rate of impairment typically observed after approximately 15 minutes that continues across the next hour.^23,38^ These findings suggest that cognitive functions associated with slower oscillations (i.e., theta band, 3-7 Hz) have a more rapid recovery than faster oscillations (i.e., beta band 13-25 Hz). It follows that sleep inertia is also characterized by different time courses in cognitive subsystem recovery (e.g., working memory task vs. simple math task).^3,23^ It may also signal a prioritization scheme of the waking brain, from high level executive functions to motor coordination; however, further research is necessary to understand the potential differences in cognitive system recovery.

### Long-range connections orchestrating local-global operations are uniquely disrupted within the brain shortly after waking

The suggested prioritization scheme in power, in addition to the delta band specificity in network changes, could suggest how this prioritization scheme is implemented in the brain. Oscillations emanating from the brain, as measured via EEG, are a consequence of short- and long-range connections within the brain that coalesce to support cognition.^39^ Slower oscillations often represent the coordination of distal regions of the cortex that modulate higher frequency oscillations within the brain.^40-45^ In other words, oscillatory activity and the associated cognitive functions may be understood as a consequence of the ever-present need and importance of global coordination of local processes.^46^ This local-global coordination of neural activity is critical to a variety of cognitive processes,^47^ is the hallmark of several diseases,^48^ and is an organizing principle of brain activity that has been suggested to be foundational even across multiple species.^39^ Within this framework, our global power results would suggest that long-range connections, as indicated by slower oscillations (delta and theta), recover more rapidly than the higher frequency bands. While global power in lower frequency bands is influenced by ensemble synaptic action across long ranges and averaged across all channels, our graph theoretic analysis is derived from the statistical dependencies between nodes, specifically estimating phase-based relationships between different channels. In other words, while global power in slower oscillations is sensitive to both amplitude and phase-based relationships aggregated across the long-range connections within the brain, our graph theoretic results are narrowly sensitive to phase-based coordination within longer connections, or the *communication structure* rather than fluctuations in synchronous neural activity. Taken together, our results display a prioritization of longer range — perhaps higher cognitive — coordinated activity after waking, while simultaneously increasing communication efficiency across them. Future research may investigate the role of higher segregation and long-range integration of networks during this process.

### Blue-enriched light serves as an intervention to mitigate neural effects of sleep inertia

We also studied an intervention condition in which participants were exposed to a blue-enriched light immediately after being awakened from SWS and throughout the one-hour testing period. Light, particularly bright, blue-enriched light, is known to have acute alerting properties when administered under conditions of sleep deprivation and is particularly effective when used during the biological night.^24,25^ EEG studies have shown that acute exposure to blue-enriched light at night during continuous wakefulness reduces delta-theta power (0.5-5.5 Hz, a biomarker of sleepiness)^49-51^ and increases alpha and high-alpha power (9.5-10.5 Hz, a biomarker of alertness).^49,51-53^

Recently, we reported the effects of light during the sleep inertia period following nocturnal awakenings. Our study showed that light modestly improved performance on a psychomotor vigilance task as well as subjective outcomes such as alertness and mood.^9^ In the current paper, we report the potential mechanisms underlying these neurobehavioral effects of light during the sleep inertia period. In contrast to previous findings during continuous wakefulness at night, we observed only modest, regional changes in power with light exposure during the sleep inertia period (see Supplemental Material). This contradiction in findings may suggest that light acts through a different mechanism during the sleep inertia period.

We observed that the significant decreases in clustering in the delta band following awakening in the control condition were attenuated with exposure to blue-enriched light. Thus, our observed effects of light on the awakening brain appear to counteract the unique hallmark of the sleep inertia period illustrated by our novel network analysis. Taken together with our lack of changes in global power with blue-enriched light exposure, this finding strengthens the specificity of the long-range communication aspect of our results. During the sleep inertia period, blue-enriched light may help to restore or protect against the dis-coordination of long-range communication within the brain. This manipulation of delta networks may represent a mechanism through which blue-enriched light exposure influences neurophysiological properties to improve alertness, mood, and performance immediately after waking.^9^ To our knowledge, no previous studies have evaluated the effect of a light stimulus on these network dynamics in different brain states; our study is the first to describe and tentatively interpret these effects.

### Methodological considerations and limitations

Although our study involved a randomized, within-subject, cross-over design with frequent testing points and high-density EEG, it is not without limitation. First, our study did not include a measure of melatonin to explore whether manipulation of its secretion was a potential contributing factor to the differences observed in the blue-enriched light condition. We assume based on prior literature, however, that melatonin was suppressed for the duration of the light exposure^49,54,55^ and, therefore, may act as a mechanism for the acute effects of light. Secondly, while a strength of our study is that we controlled the sleep stage at awakening, this was traded for potential changes in circadian sleep pressure between the two awakenings. The order of condition was randomized to limit any differences, and the mean time between awakenings was 90 minutes,^9^ so we do not expect large differences from this design. Further, the likelihood of a circadian effect in the current study is low as we only observed the reduction in clustering and path length immediately after awakening, with a rapid return to baseline levels, which is not characteristic of a circadian effect. We acknowledge that we are unable to directly disentangle the relative contributions from, or effect of light on, the three sleep processes (homeostatic, circadian, inertia) in the current study. However, our first testing point (T1 at 2 minutes post-awakening) is a robust proxy for sleep inertia. Importantly, we have, for the first time, demonstrated the neurophysiological profile of the waking brain and the effect of blue-enriched light exposure following awakening from SWS during a nocturnal sleep episode, which is a common scenario for on-call and emergency service workers.

### Conclusions

The current study extends prior research investigating the waking brain by building a more comprehensive description of the neurophysiological profile of the brain following awakening from SWS at night. Our results suggest that long-range network communication within the brain is crucial to the waking process and, further, that the brain may prioritize these long-range connections, adding to the evolutionary importance of the coordination of local and global activity within the brain. Moreover, the addition of a within-subject assessment of the effects of blue-enriched light exposure provides more insight into our understanding of sleep inertia, adds a causal aspect to our findings, and suggests how we might mitigate its effects to improve alertness and performance in safety-critical scenarios.

## Materials and Methods

### Participants

Twelve healthy young adults participated in the study having met the following inclusion criteria: healthy (General Health Screening Questionnaire, personal physician’s permission to participate, approval from onsite physician upon review of urinalysis and blood work screening); normal sleepers (Pittsburgh Sleep Quality Index ≤5; no self-reported sleep problems; habitual sleep of 7-9 hours); no shiftwork or travel >3 time zones in the past 3 months (self-report); free of illicit substances and nicotine (urine toxicology screen); and free of alcohol during the study period (breathalyzer). One participant’s dataset was incomplete, therefore, results presented here reflect a sample population of *n* = 11.

*A priori* power calculations were based on anticipated changes in our primary outcome measure, psychomotor vigilance test (PVT) performance. Based on previous studies in similar populations, we expected that the effect size for the change in PVT during sleep inertia would be approximately 0.75. Using these assumptions, we estimated that we would need 10 participants to detect a change in performance with 80% power at an alpha level of .05.

### Procedure

The results presented here come from a two-week study. Here we report a one-week within-subject, cross-over intervention study with the presentation order of intervention randomized by sex.

Participants were required to follow a fixed sleep-wake schedule based on habitual sleep timing for the six nights leading up to the in-laboratory visit (see Figure 4). Following a night of at-home sleep restriction (5 hours), participants were brought into the sound attenuated, light- and temperature-controlled laboratory for pre-sleep procedures that included task familiarization, electrode set up, and baseline tests prior to overnight observation and testing.

**Figure 4.**
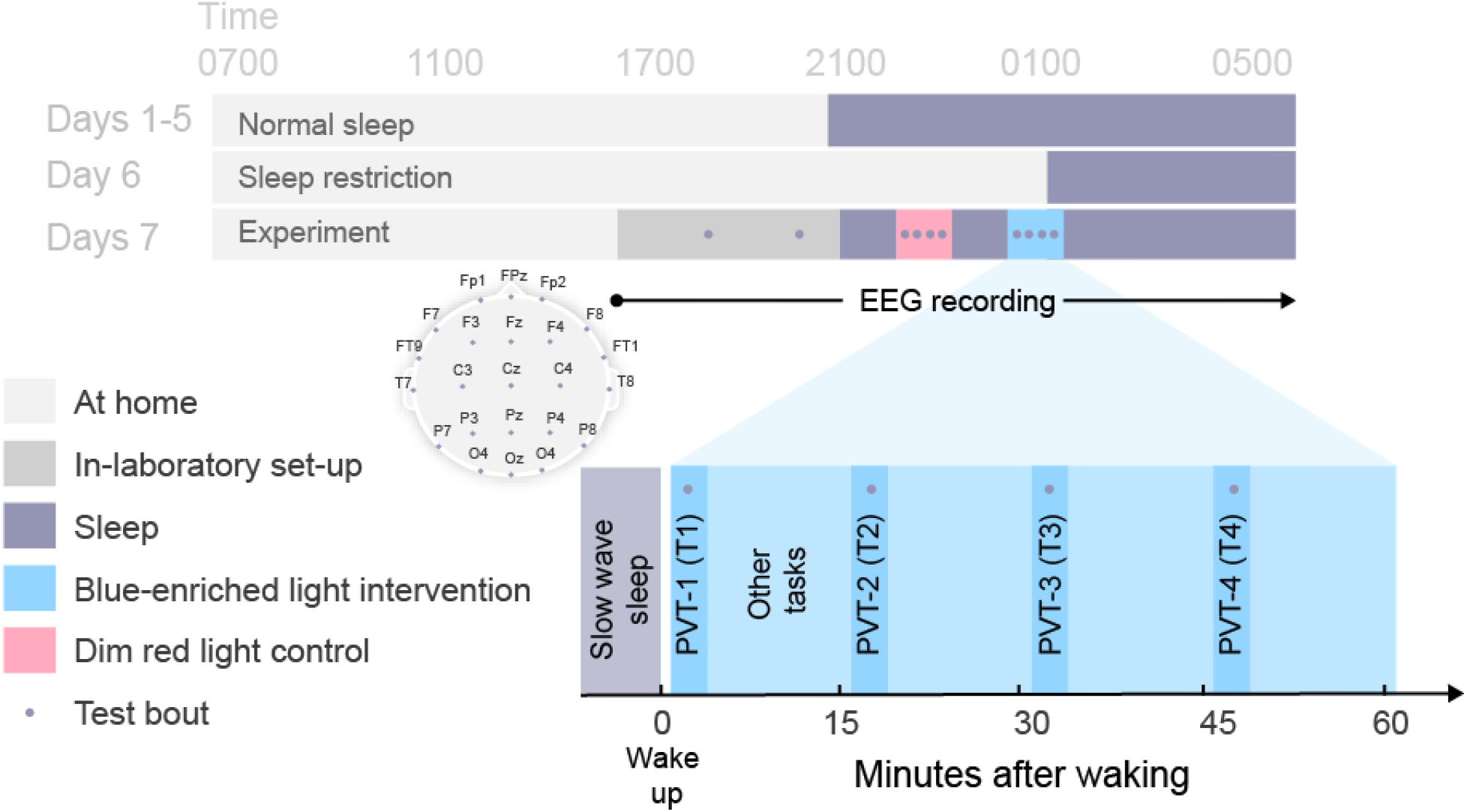
Protocol schematic. Light gray shading indicates wakefulness during the at-home portion of study. Dark gray shading indicates in-laboratory pre-sleep activities including baseline testing (•). Black shading indicates sleep opportunities (<0.3 lux). Blue and red shading indicate intervention and control sleep inertia testing periods, respectively. Inset shows electrode montage and post-awakening test bouts. Clock times shown are approximate and varied depending on habitual sleep-wake times and appearance of slow wave sleep periods.

At the participant’s habitual bedtime, all lights were turned off (<0.3 lux) and the participant was instructed to sleep. Electroencephalography (EEG) was monitored during the sleep period to identify slow wave sleep (SWS) stages (Stage 3 and 4).^56^ Participants were awakened after a minimum of five consecutive minutes of SWS. Immediately upon awakening, a dim, red ambient light was illuminated in the room. In the intervention condition only, at one minute post-awakening, a blue-enriched light was illuminated and remained on for the hour of testing. The dim, red, ambient light remained on during the testing period in both conditions. A five-minute PVT was performed four times (T1, T2, T3, and T4 at +2, +17, +32, and +47 minutes after the awakening, respectively), during which EEG was recorded. At the end of the testing period, all lights were turned off and the participant was instructed to return to sleep. EEG was monitored again to identify the next period of five consecutive minutes of SWS, at which time the participant was awoken again and exposed to the opposite condition (blue-enriched light or control). Following the second testing period, all lights were turned off and the participant was instructed to sleep until they were awakened at their habitual wake time.

### Intervention

A 12” x 24” canvas of blue-enriched light-emitting diodes (LEDs) (Circadian Positioning Systems, Inc., Newport, RI) was positioned at 15 degrees to the horizontal angle of gaze and approximately 56 cm away from the participant. Light levels during the intervention and control conditions were confirmed via Spectroradiometer ILT950 (International Lighting Technologies, Peabody, MA). Illuminance, irradiance, equivalent daylight (D65) illuminance (EDI), and peak spectra during the intervention were 242.77 lux, 0.95 W/m^2^, 338.03 melanopic lux, and 456 nm, respectively, measured at the angle of gaze. An ambient dim, red light served as the control (0.26 lux, 0.00 W/m^2^, 0.10 melanopic lux, 714 nm).

### EEG Analysis

#### Pre-processing

High density EEG was recorded during the baseline and post-awakening testing periods using BrainVision 32-channel caps with sintered Ag/AgCI electrodes (Brain Products GmbH, Munich, Germany) and BrainVision Recorder software (Brain Products GmbH, Munich, Germany). Additional electrodes included bipolar horizontal electrooculogram (EOG: left/right placed on the outer canthus of the eye 1 cm above and below the horizon, respectively), and sub-mental electromyogram (EMG), for standard monitoring with sleep periods. For visualization while the experimenter monitored the EEG, a 70 Hz high-pass filter in conjunction with a notch filter at 60 Hz was used online so that the experimenter could easily determine whether the participant was in SWS and primed for an awakening. After the recording sessions, the raw, unfiltered EEG recordings were then subjected to a thorough artifact editing scheme offline. After a preliminary filtering of the raw EEG data, using a third order zero-phase bandpass Butterworth filter (0.5 Hz -50 Hz) in EEGlab^57^. the EEG data were subjected to Artifact Subspace Reconstruction (ASR).^58-60^ This method removes extremes in data using a time-evolving blind source separation method; importantly, this method has been shown to be particularly resilient to artifact encountered in real-world scenarios.^61^ To deploy ASR on the dataset, we first created a “clean” reference signal from each participant’s EEG data by concatenating EEG segments that were at least 1000 ms long with amplitude below 100 μV, most likely not contaminated by artifact due to muscle activity following the awakening. Following the creation of the reference signal, ASR was then used to clean the EEG that contained large fluctuations greater than five standard deviations beyond the reference signal (in 500 ms chunks). Finally, EEG data from the beginning to the end of each PVT (approximately 5 minutes each) was identified and then filtered via a third order zero-phase bandpass Butterworth filter within the frequency bands of interest (delta: 1-3 Hz, theta: 4-7 Hz, alpha: 8-12 Hz, beta: 15-25Hz). This produced four sets of continuous time courses for each of the time segments following the awakening (T1, T2, T3, T4) in addition to the baseline pre-sleep time period (BL) for each participant.

#### Global power spectral density

Power spectral density (PSD) was estimated using a standard approach of Welch’s average modified periodogram method of spectral estimation^62^ in Matlab (Mathworks, Inc.). The log-transformed PSD values were then standardized for each electrode before analysis by mean-centering each channel across frequency. For the sake of simplicity, hereafter, we use the term ‘power’ to represent the standardized PSD values and global power is the average electrode standardized PSD across the scalp.

#### Network Connectivity

To estimate the functional network connectivity between EEG sensors, we computed the pairwise Weighted Phase Lag Index (wPLI), which is known to be highly sensitive to linear and nonlinear interactions;^63^ it is well-established that phase-based measurements of connectivity are less susceptible to nuisance artifacts.^64^ Within a participant, we calculated the wPLI matrix for all the time points and frequency bands by using the band-wise filtered EEG activity for the entire 5-minute trial epoch. This produced one matrix for T1, T2, T3, T4, and the baseline pre-sleep time period.

#### Network Analysis

Three network measurements were estimated on the wPLI matrices, including clustering coefficient, path length, and (in supplemental) degree. These common metrics have been used to describe properties of many complex systems including a variety of biological, social, and other phenomena^15,65,66^ and are often evoked when describing small-world phenomena.^65^ Here, we use clustering coefficient and path length to describe network changes related to small-worldness/randomness of the network and degree as a visualization of changes in network connectivity.

Specifically, clustering coefficient (C) estimates the tendency of a node’s neighbors within a network to also be linked, and may be mathematically described from a connectivity matrix, *W* (here estimated via wPLI).^67,68^

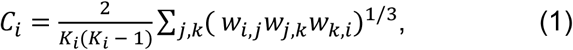

where *w*_*i,j*_ represents an element of the connectivity matrix *W* implying the strength of connectivity between nodes *i* and *j. C*_*i*_ and *K*_*i*_ represent the clustering coefficient and degree for node *i*. Degree of a node is defined as the sum of all the edge weights connected to it and is a general representation of a node’s connectivity across the network:

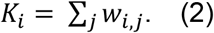

*Path length*, on the other hand, estimates the number of edges, on average, that must be traversed to connect any two nodes within a network. If *d(i, j)* represents the shortest path through edges between nodes *i* and *j*, Path length *λ* is given by:^68^

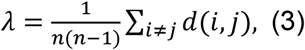

where, *n* is the total number of nodes. In general, the distance between two nodes with strong connectivity is lower than the distance between nodes with relatively weaker connectivity. Therefore, to estimate *d(i, j)*, we used (1-*w*_*i,j*_) and then the path length was calculated using Brain Connectivity Toolbox function *charpath*.*m*.

#### Analysis & Statistics

Our primary analysis explored the network configuration of the awakening brain under control conditions. Our secondary analysis explored the impact of blue-enriched light on this network profile by comparing conditions (control and light). Much of our analyses relied on statistical comparisons between the pre-sleep baseline period and each of the post-sleep test bouts. Thus, for each frequency and metric of power or connectivity and condition (e.g., control, blue-enriched light), the average metric for each individual and time period were submitted to paired *t*-tests in Matlab (Mathworks, Inc.). Where appropriate for multiple comparisons, false discovery rate was used with a *q* threshold set to .05.

#### Visualization

All figures, boxplots, and topographic plots were created in Matlab (Mathworks, Inc.) with common core functionality from Matlab and some additional functions with EEGlab.^57^ Then, they were imported to Adobe Illustrator (version 25.3) and combined into panels for visualization.

## Acknowledgements

This work was supported in part by the Naval Medical Research Center’s Naval Advanced Medical Development Program (MIPR N3239820WXHN007) and the NASA Airspace Operations and Safety Program, System-Wide Safety. This work was also supported in part through mission funding from the US DEVCOM Army Research Laboratory (ARL). K.B. also acknowledges support from ARL through Cooperative Agreement Number W911NF-16-2-0158. The views and conclusions contained in this document are those of the authors and should not be interpreted as representing the official policies, either expressed or implied, of the ARL or the US Government. The authors would also like to thank Aditi Periyannan for her help with figure design and layout. The authors would also like to thank all the participants who volunteered their time for this study.

