## Supplemental Material for "Reconfigurations in brain networks upon awakening from slow wave sleep: Interventions and implications in neural communication"

**Supplemental Materials**

Table S1. Participant Demographics (*n* = 11)

| | | | |
|---|---|---|---|
| | *n* (%) | | |
| Female | 6 (55%) | | |
| | Mean | SD | Range |
| At-home sleep duration (h) | | | |
| Nights 1-5 | 7.2 | 0.6 | 6.4 – 8.2 |
| Night 6 | 4.4 | 0.3 | 3.8 – 5.1 |
| Questionnaires | | | |
| PSQI | 2.1 | 1.2 | 0 – 4 |
| FSS | 26.7 | 7.0 | 16 – 39 |
| MEQ | 55.5 | 6.9 | 45 – 64 |

*Note*: Sleep variables estimated by actigraphy: sleep duration = sleep period minus wake after sleep onset. Range and SD are based on participant means. SD = standard deviation; h = hour; PSQI = Pittsburgh Sleep Quality Index; FSS = Fatigue Severity Scale; MEQ = Morningness-Eveningness Questionnaire.

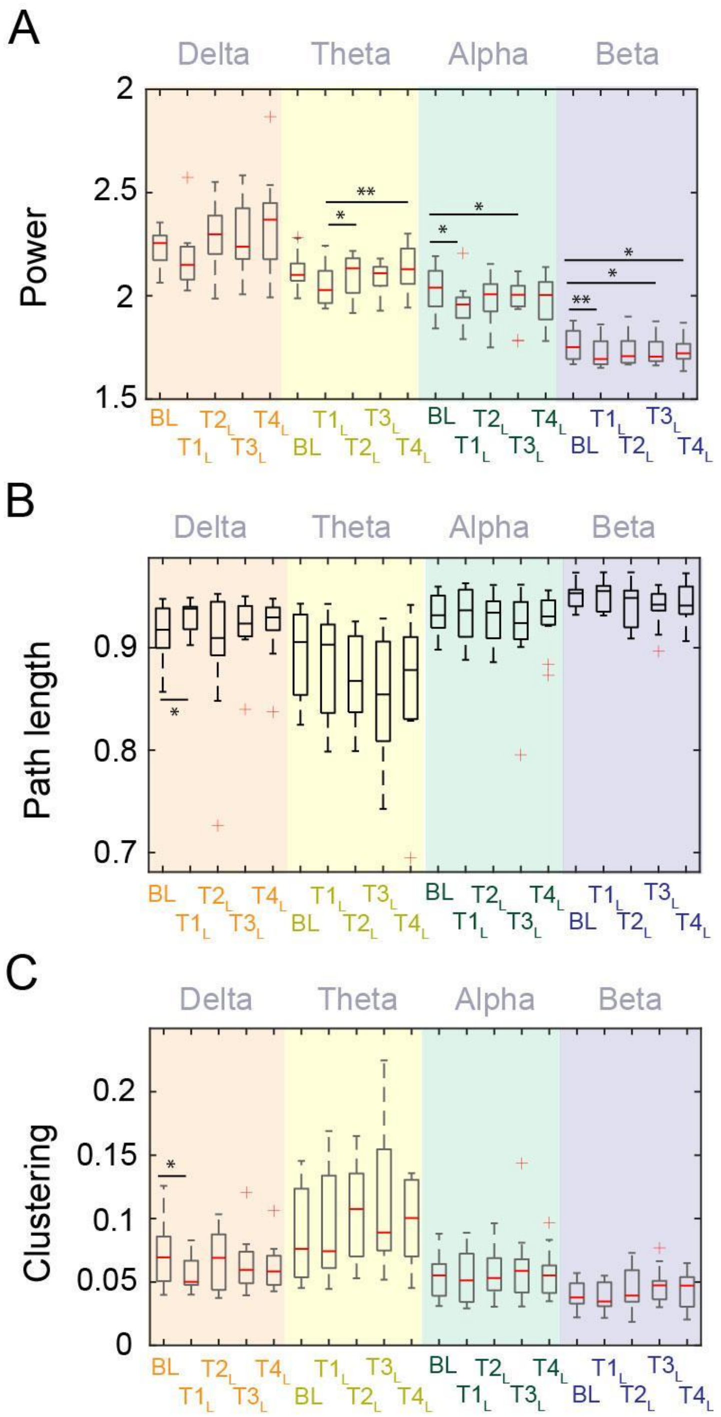


Figure S1. (A) Comparison for delta power, and (B) - (C) brain network properties across test bouts for each frequency band in the light intervention condition (blue-enriched light). BL = baseline, $T\#_L$ = Test bout # during the light condition. *$p < .05$ and **$p < .01$.

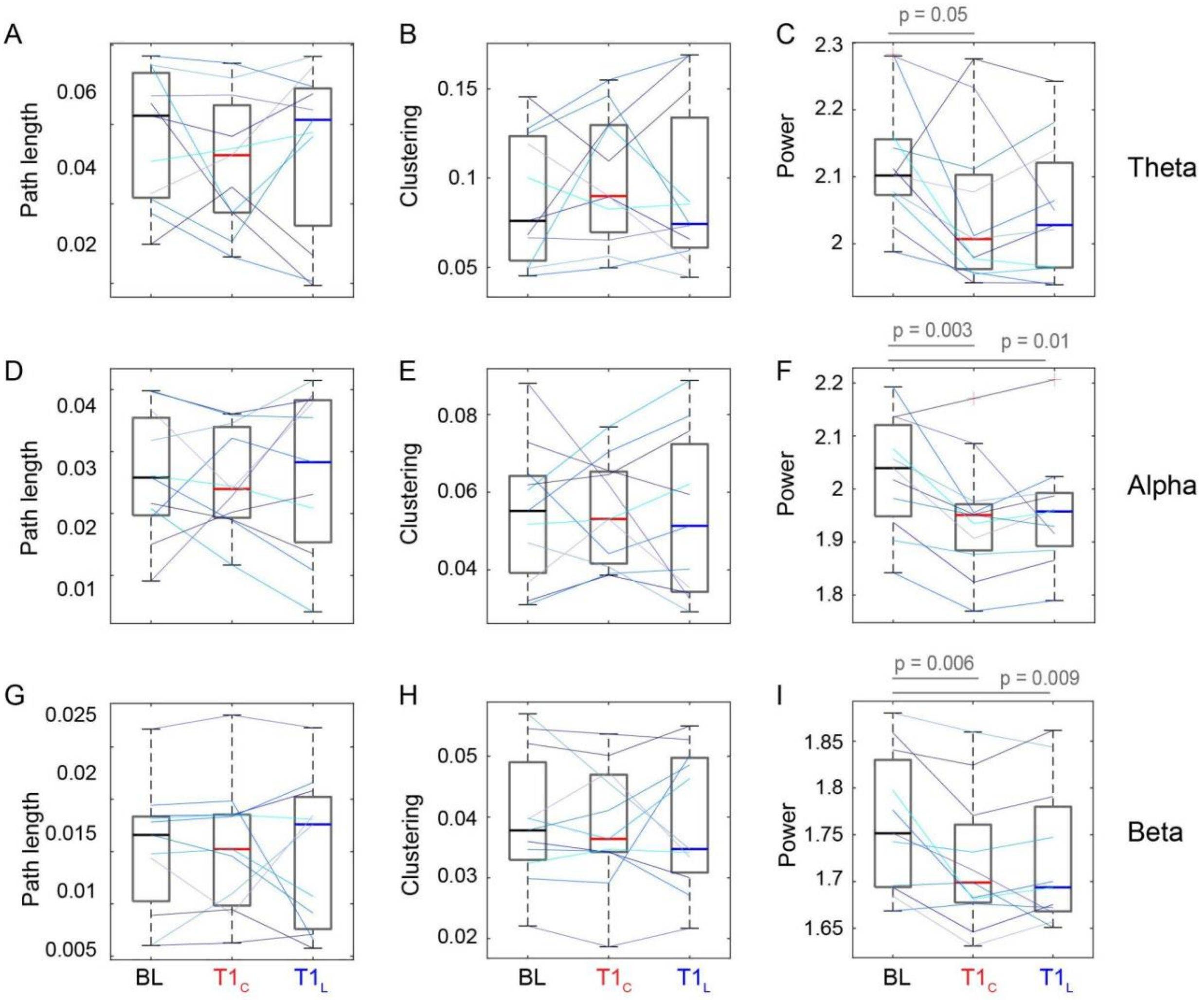


Figure S2. (A) - (B) Brain network properties comparing pre-sleep baseline (BL), control at T1 ($T1_C$), and light at T1 ($T1_L$) for different frequency bands; (C) Similar comparison for power. Colored lines represent individual participants.

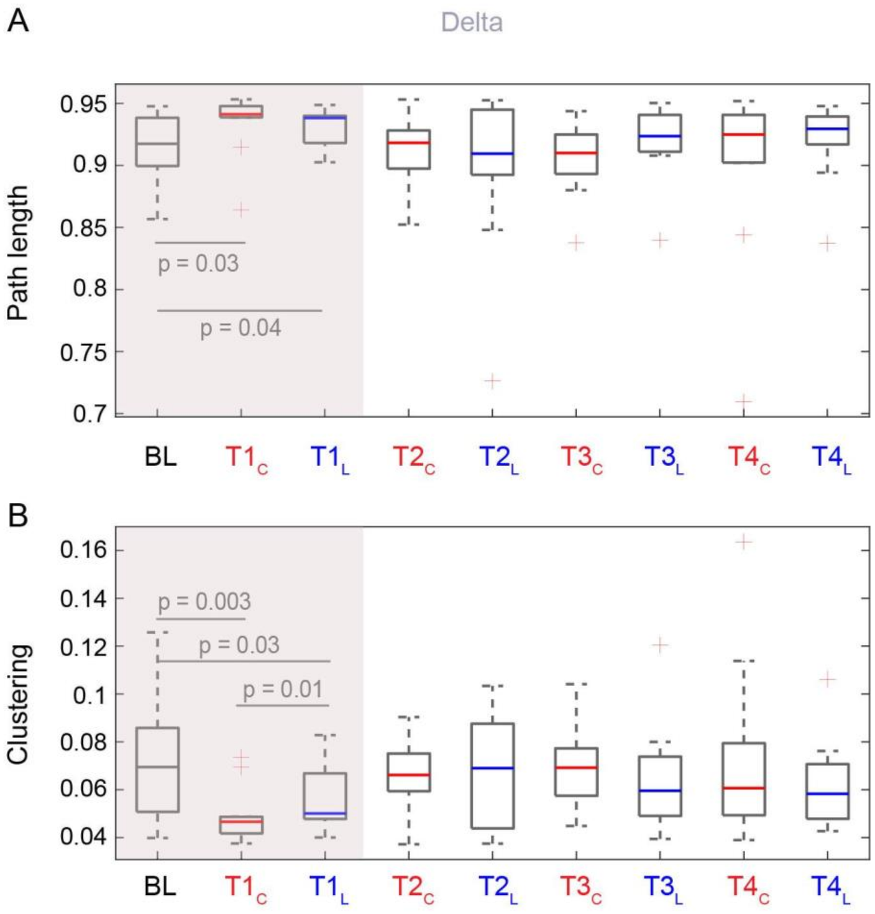


Figure S3. Brain network properties comparing pre-sleep baseline (BL), control, and light at different test bouts (T1 to T4) for the delta frequency band. The shaded area is reproduced from Figure 3. Here, significant (< .05) *p*-values are indicated only for the *t*-test comparisons with the baseline and between control and light within each test bout.

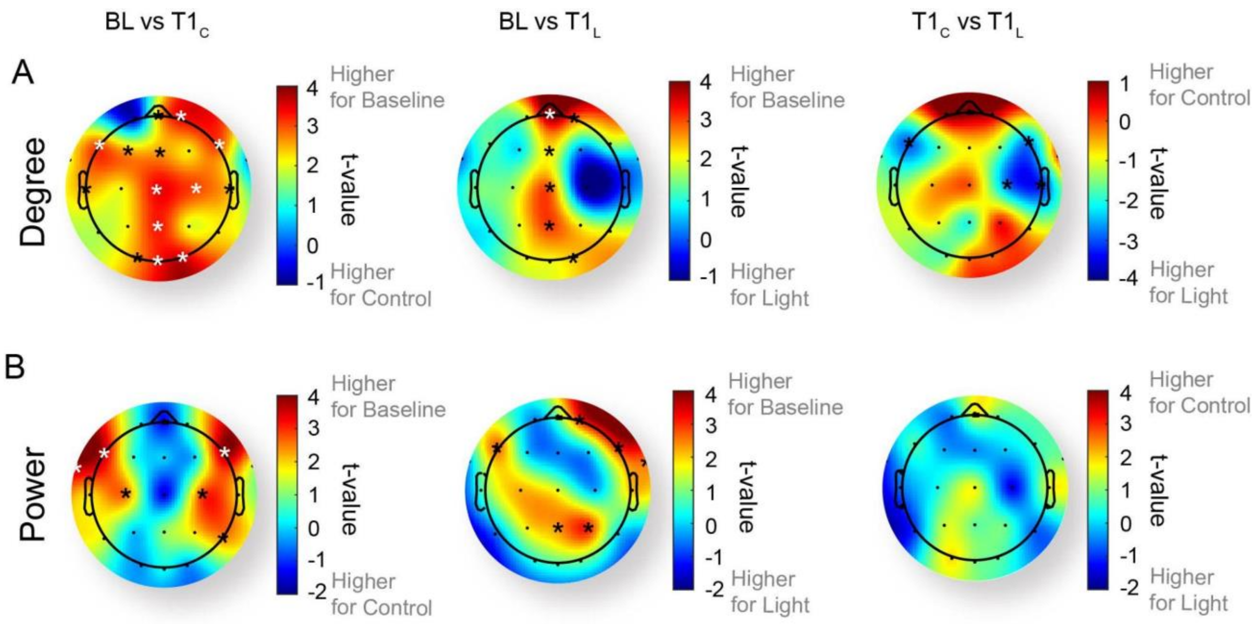


Figure S4. Change in (A) Degree and (B) Power between baseline (BL) and control at T1 ($T1_C$), baseline and light at T1 ($T1_L$), and control and light at T1 across scalp regions. White asterisks represent electrodes with significance ($q < .05$) when corrected for multiple comparisons; black asterisks represent uncorrected significance ($p < .05$).
